# Phagolysosomes break down the membrane of a non-apoptotic corpse independent of macroautophagy

**DOI:** 10.1101/2024.06.19.599770

**Authors:** Shruti Kolli, Cassidy J. Kline, Kimya M. Rad, Ann M. Wehman

**Author notes:** Equal contribution (Shruti Kolli, Cassidy J. Kline).

## Abstract

Cell corpses must be cleared in an efficient manner to maintain tissue homeostasis and regulate immune responses. Ubiquitin-like Atg8/LC3 family proteins promote the degradation of membranes and internal cargo during both macroautophagy and corpse clearance, raising the question how macroautophagy contributes to corpse clearance. Studying the clearance of non-apoptotic dying polar bodies in *Caenorhabditis elegans* embryos, we show that the LC3 ortholog LGG-2 is enriched in the polar body phagolysosome independent of membrane association or autophagosome formation. We demonstrate that ATG-16.1 and ATG-16.2, which promote membrane association of lipidated Atg8/LC3 proteins, redundantly promote polar body membrane breakdown in phagolysosomes independent of their role in macroautophagy. We also show that the lipid scramblase ATG-9 is needed for autophagosome formation in early embryos but is dispensable for timely polar body membrane breakdown or protein cargo degradation. These findings demonstrate that macroautophagy is not required to promote polar body degradation, in contrast to recent findings with apoptotic corpse clearance in *C. elegans* embryos. Determining how membrane association of Atg8/LC3 promotes the breakdown of different types of cell corpses in distinct cell types or metabolic states is likely to give insights into the mechanisms of immunoregulation during normal development, physiology, and disease.

## Introduction

Throughout an organism’s life, cells die and must be degraded in a contained manner. Phagocytosis is an essential process by which dying cells, cellular debris, and pathogens are cleared from the extracellular space and degraded inside phagolysosomes. Disrupting engulfment and intracellular degradation of dying cells can cause inflammation and lead to the development of autoimmune diseases (1). For the contents of dying cells to be degraded, their plasma membrane must be broken down inside the phagolysosome, however how this is regulated remains unclear.

Previous work has implicated lipidation of the ubiquitin-like Atg8/LC3 family proteins in promoting phagolysosomal degradation of cell corpse membranes and their cargo (2). Depleting ATG-7, part of the Atg8/LC3 lipid conjugation machinery in *C. elegans*, delayed breakdown of the membrane of a non-apoptotic cell corpse and delayed disappearance of its contents in phagolysosomes, suggesting that lipidation of Atg8/LC3 family proteins promotes corpse degradation. Atg8/LC3 family proteins are better known for their roles during macroautophagy, when intracellular cargos are enclosed in a double membrane for degradation (3). Lipidation of Atg8/LC3 family proteins promotes breakdown of the inner membrane of autophagosomes during macroautophagy in mammalian cells (4), suggesting that lipidated Atg8/LC3 proteins have a conserved role in promoting the breakdown of internal membranes inside phagolysosomes and autolysosomes.

Several studies have found that when autophagosome formation or maturation was disrupted in *C. elegans* embryos, degradation of apoptotic corpses was significantly delayed within phagolysosomes (5–7). Recently, it was shown that autophagosome fusion with phagosomes promotes apoptotic corpse clearance mid-embryogenesis (8). Autophagosome puncta of fluorescent Atg8/LC3 family proteins LGG-1 and LGG-2, as well as a fluorescent transmembrane autophagy protein ATG-9, were observed next to phagosomes containing apoptotic cargo. As the bright LGG-1, LGG-2, and ATG-9 puncta disappeared, the markers were observed inside phagolysosomes, consistent with autophagosome fusion with the phagolysosome to transfer the internal membrane of the autophagosome into the phagolysosome. In addition, these autophagosome reporters were dispersed in the phagolysosome lumen, indicating prior breakdown of the corpse plasma membrane within the phagolysosome (8). These results suggest that autophagosome fusion contributes to the degradation of apoptotic cargo in phagolysosomes.

LGG-1 and LGG-2 reporters were also observed inside a non-apoptotic phagolysosome after corpse membrane breakdown (2), raising the question whether autophagosomes play a role in the early embryonic clearance of the second polar body by undifferentiated blastomeres. Polar bodies are born during meiosis and show TUNEL labeling independent of apoptotic regulators (9), as well as a loss of membrane integrity commonly associated with necrosis (2). *C. elegans* polar bodies are well-suited to study the dynamics of phagocytosis and cargo resolution, given the stereotyped timing of internalization by large embryonic blastomeres and membrane breakdown inside the phagolysosome (Fig 1). However, polar body membrane breakdown was not delayed when the ATG14 ortholog EPG-8 was deleted (2). ATG14 family proteins are thought to act as part of the PI3Kinase complex during canonical autophagy only (10), suggesting that the localization of Atg8/LC3 family proteins inside polar body phagolysosomes was independent of autophagosomes and canonical autophagy. Given the conflicting results between the role of autophagosomes in apoptotic and non-apoptotic cell clearance during different stages of *C. elegans* embryogenesis, we revisited whether canonical autophagy plays a role in breakdown of the polar body membrane using proteins that act upstream in the macroautophagy pathway, specifically homologs of Atg9 or ATG16L1.

**Fig 1.**
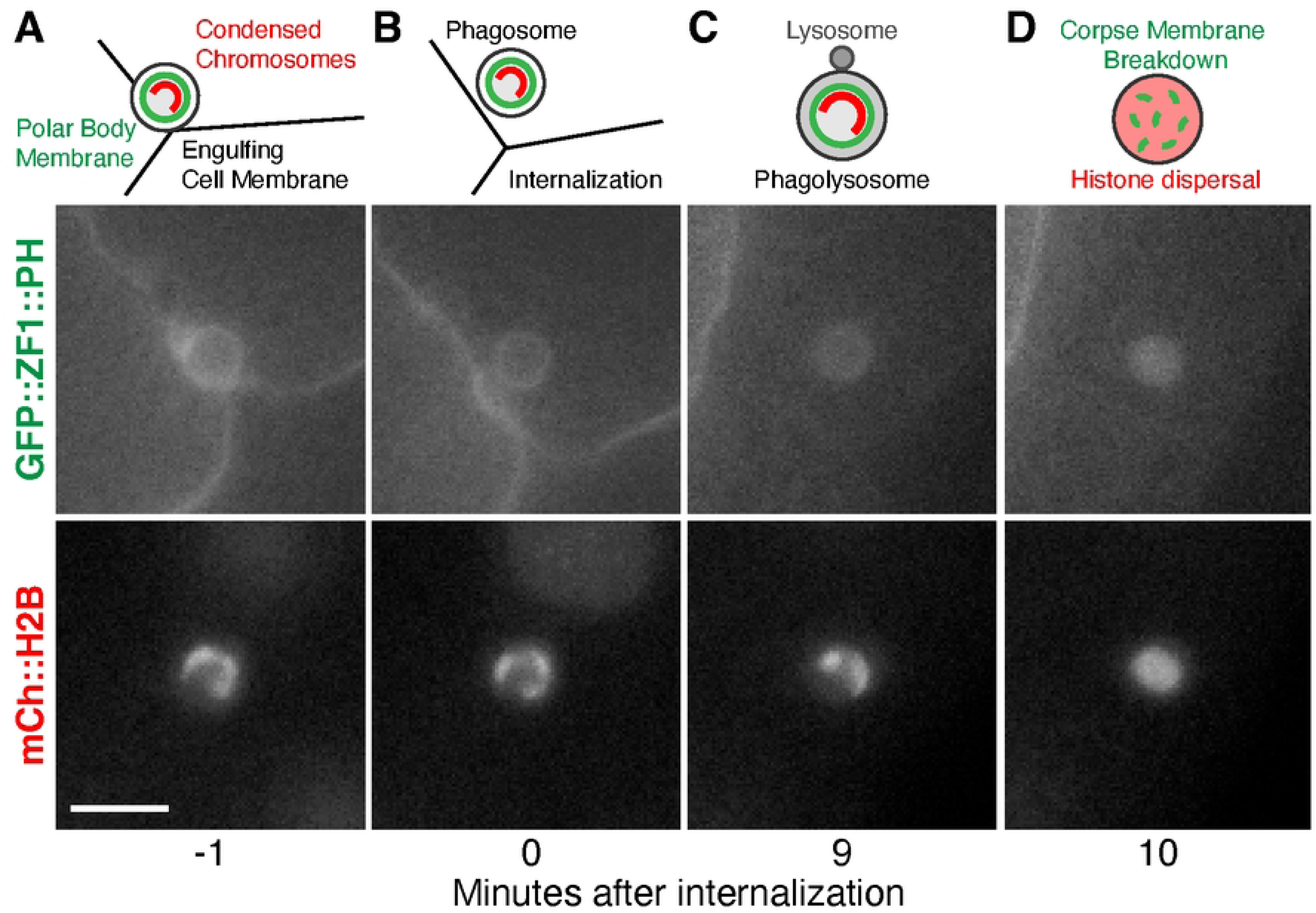
Engulfment of the second polar body by embryonic blastomeres and corpse membrane breakdown inside the phagolysosome. A) Diagram and still images of the second polar body next to three embryonic blastomeres expressing GFP::ZF1::PH to label plasma membranes and mCh::H2B to label histones. Scale bar is 5 µm. B). After engulfment, the polar body is found inside a phagosome within the cytosol of one blastomere. C) The phagosome fuses with lysosomes (L) to become a phagolysosome, but the plasma membrane of the polar body initially remains intact, and the polar body chromosomes remain condensed. D). After phagolysosomal enzymes disrupt the plasma membrane of the second polar body, both the GFP::ZF1::PH membrane and mCh::H2B histone reporters disperse within the phagolysosome lumen.

The lipid scramblase Atg9 acts early in autophagosome biogenesis, during the expansion of phagophores to engulf their intracellular cargo (11). The lipid transfer protein Atg2 transfers lipids to one leaflet and Atg9 acts as a lipid channel to distribute phospholipids across the leaflets of the phagophore bilayer, enabling the growth of isolation membranes.

Apoptotic corpses accumulate in *atg-9* mutants (7), and *C. elegans* ATG-9 was recently shown to localize inside phagolysosomes clearing apoptotic corpses after autophagosome fusion (8). However, it is unknown whether ATG-9 plays a role during the phagolysosomal breakdown of non-apoptotic polar bodies.

Mammalian ATG16L1 targets LC3 lipidation to specific membranes (12), using different domains of the protein during canonical and non-canonical autophagy (13, 14). The N-terminal domain of ATG16L1 binds to the ATG5-ATG12 heterodimer and promotes LC3 lipidation to autophagosome, phagosome, and endosome membranes. The central coiled-coil domain (CCD) binds WIPI2 or FIP200 and is required for macroautophagy, while the C-terminal WD40 domain is required for non-canonical roles of Atg8/LC3 lipidation, such as LC3-associated phagocytosis (LAP) or conjugation of Atg8 to single membranes (CASM). *C. elegans* has two ATG16L1 orthologs, ATG-16.1 and ATG-16.2, which function redundantly to localize lipidated Atg8/LC3 family proteins LGG-1 and LGG-2 to autophagosomes (15).

ATG-16.2 has a more important role during autophagy than ATG-16.1, with lipidation of Atg8/LC3 family proteins occurring in *atg-16* mutants, but LGG-1 and LGG-2 fail to localize to autophagosomes. Both ATG-16 orthologs promote autophagic degradation in embryos (15), but a role during LAP or CASM has not been examined. As ATG16L1 recruits Atg8/LC3 to both double-membrane autophagosomes and single-membrane phagosomes (13, 14), we used *C. elegans atg-16* mutants to test whether Atg8/LC3 localization to autophagosomes or other membranes promotes breakdown of the polar body membrane inside phagolysosomes.

In this study, we confirm that ATG-9 and ATG-16.2 are required for autophagosome biogenesis in *C. elegans* early embryos. In the absence of autophagosomes, we find no delay in corpse membrane breakdown, no disruption in LGG-2 localization inside the polar body phagolysosome, and no delay in degradation of a corpse cargo protein. We also confirm the role of membrane association of Atg8/LC3 by demonstrating that ATG-16.1 and ATG-16.2 redundantly promote polar body membrane breakdown. This suggests that autophagosomes are not required for phagolysosome maturation in early embryos, and the association of Atg8/LC3 with a non-autophagic membrane is important for the phagocytic clearance of a non-apoptotic corpse.

## Materials and Methods

### Worm maintenance

*Caenorhabditis elegans* strains (Table S1) were maintained at room temperature (22-24°C) on nematode growth media (NGM) plates seeded with OP50 bacteria, according to standard procedures (16). *C. elegans* grown for semi-quantitative PCR were maintained on peptone-rich plates seeded with NA22 bacteria at 18.5°C.

### Genotyping

DNA from lysed worms was amplified using OneTaq polymerase (New England BioLabs) and the primers listed in Table S2 that were designed based on sequences in WormBase (17). MboI (New England BioLabs) was used to detect the point mutation in *atg-9(bp564)* and HinfI (New England BioLabs) was used to detect the point mutations in atg*-16.1(gk668615[Q356*])* and *atg-16.2(gk145022[W253*])*.

### Transgenesis

The CTPD fragment from pDonr221-oma-1 C-term (18) was amplified using the primers mex-5p oma-1(219) F and mCh oma-1(378) R (Table S2) and then cloned into pVIG57 (gift of Vincent Galy) using a HiFi reaction (New England Biolabs). The resulting plasmid pCFJ150_Pmex-5:CTPD::mCh::LGG-2 was injected for MosSCI (19) into strain COP93 by InVivo Biosystems. The inserted transgene was homozygosed to generate strain WEH751.

### Microscopy

Gravid hermaphrodites were dissected in egg salts (94 mM NaCl, 32 mM KCl, 2.8 mM MgCl_2_, 2.8 mM CaCl_2_, 2 mM HEPES, pH 7.5) to isolate embryos and mounted on a 4% agarose pad for live imaging. Z-stacks were collected using a Zeiss Axio Observer 7 inverted microscope with Plan-APO 40X 1.4 NA oil objective with Excelitas Technologies X-Cite 120LED Boost illumination, and Hamamatsu ORCA-Fusion sCMOS camera controlled by 3i SlideBook6 software. Time-lapse imaging was acquired sequentially for mCherry and GFP every minute.

### mCh::LGG-2 autophagosome count

Bright mCherry puncta in the cytosol of 8- to 15-cell embryos were counted in SlideBook (3i) or using the cell counter function in FIJI (20). Larger and dimmer LGG-2-positive puncta flanking dividing nuclei appeared to be centrosomes and were not included in the autophagosome count. Large clusters of puncta were challenging to distinguish and omitted from autophagosome counts in 1-cell embryos in Fig 7C.

### Qualitative colocalization analysis

Images of live 8- to 15-cell embryos were analyzed in SlideBook (3i) for whether mCh::LGG-2 levels appeared higher at the second polar body than in the neighboring cytoplasm. Polar bodies and membrane breakdown were identified using GFP::H2B.

### Fluorescence intensity measurements

The mean intensity of a circle of 1.59 µm diameter (10 pixel) for polar bodies and 3.98 µm (25 pixel) for AB or P1 blastomere nuclei was measured in the mCh::LGG-2 channel on the polar body or nuclei and three neighboring regions of the cytoplasm using FIJI. The results in Fig 5G, 6C-D, and 7F are reported after subtraction of the average cytoplasm mean from the polar body or blastomere nuclei mean.

### Polar body internalization

Internalization was determined using the GFP::PH membrane marker as the timepoint in which the second polar body was fully engulfed by a blastomere, as previously described (21). Cell stages were identified as the beginning of each furrow ingression.

### Polar body membrane breakdown

Polar body membrane breakdown was scored indirectly using mCherry histone markers, based on established protocol (21). Membrane breakdown was the first timepoint in which the histones dispersed to fill the entire polar body phagosome.

### Polar body clearance

Polar bodies were tracked in time-lapse series using the ZF1::mCh::H2B marker. Clearance was determined as the first timepoint in which the largest polar body fragment was indistinguishable from background noise.

### Semi-quantitative RT-PCR analysis

Frozen worm pellets were vortexed with glass beads (Sigma-Aldrich) and Trizol (Zymo Research). RNA purification was performed using the Direct-zol RNA Miniprep Plus kit (Zymo Research) and reverse transcription was performed using RNA to cDNA EcoDry Premix (TaKaRa). 100ng cDNA was amplified through a 30-cycle PCR protocol using OneTaq polymerase (New England BioLabs) and the primers indicated in Table S2. SYBR Safe fluorescence (Invitrogen) was photographed using an iPhone 13 (Apple). The green fluorescence intensity was measured for each band and on empty regions of the gel using FIJI. The background fluorescence was subtracted from the band fluorescence and then normalized to the average intensity of the wild-type N2 bands.

### Image processing

Single z planes are shown, except for Fig 2A-E, where 26 z planes at 0.77 µm steps were projected (maximum intensity), Fig 3A, where 8 z planes at 1.5 µm steps were projected (maximum intensity) and Fig 6A-B, where 3 z planes at 1.5 µm steps were projected for mCh::LGG-2 (maximum intensity) using Slidebook. Images were cropped, rotated, and their brightness was adjusted in Adobe Photoshop.

**Fig 2.**
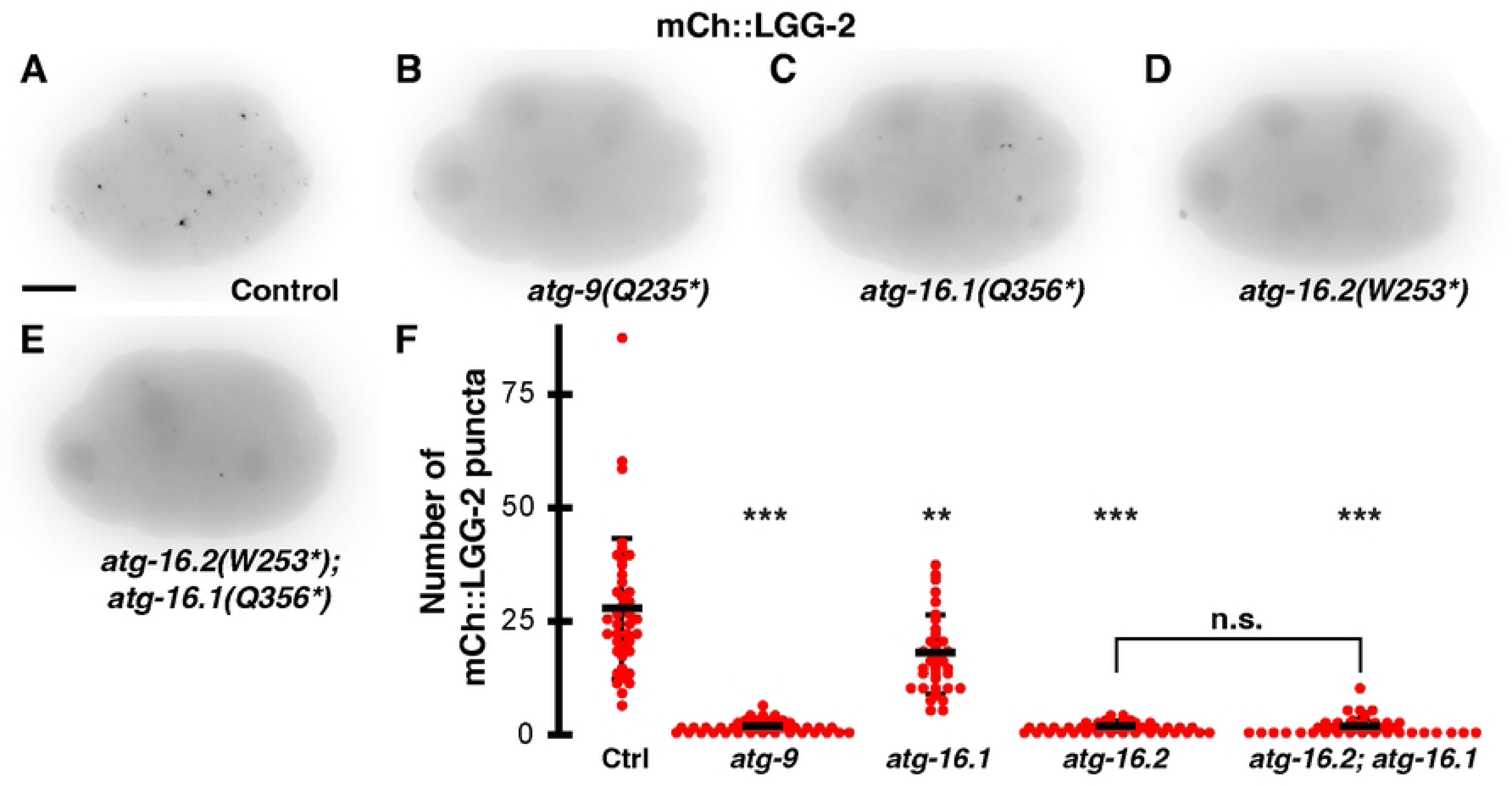
ATG-9 and ATG-16.2 are required for LGG-2-positive autophagosome formation in early embryos. (A-E) Inverted images of 8-cell embryos expressing mCh::LGG-2. Puncta were rarely observed in *atg-9* (B), *atg-16.2* (D), or *atg-16.2; atg-16.1* (E) mutant embryos compared to control embryos (A). Puncta were reduced in *atg-16.1* mutant embryos (C). Scale bar is 10 µm. (F) Control embryos averaged 27±15 mCh::LGG-2 puncta per 8- to 15-cell embryo (n=40). The *atg-9(Q325*)* and *atg-16.2(W253*)* mutants averaged significantly fewer, 1±1 mCh::LGG-2 puncta per embryo (n=41 and 40). The reduction in mCh::LGG-2 puncta was more moderate in *atg-16.1(Q356*)* mutants (17±9, n=39), but double mutant *atg-16.2(W253*); atg-16.1(Q356*)* embryos averaged 1±2 mCh::LGG-2 puncta (n=39). Data are presented as mean ± std dev. One-tailed t-test with a Bonferroni correction for 4 comparisons: **p<0.01, ***p<0.001. n.s. = not significant p>0.1.

**Fig 3.**
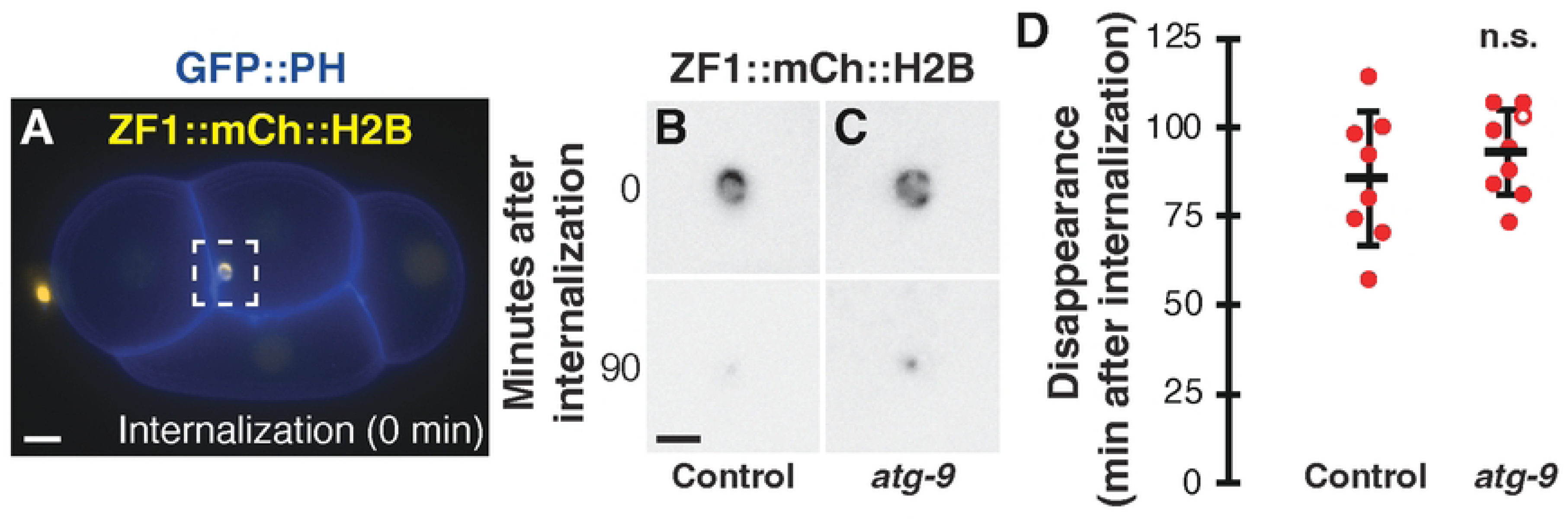
Macroautophagy does not promote degradation of polar body protein cargo. (A) A 4-cell control embryo expressing ZF1::mCh::H2B (yellow) and GFP::PH (blue) internalizing the dying polar body in a phagosome (time 0). Scale bar is 5 μm. (B-C) Inverted images of ZF1::mCh::H2B in the polar body phagosome at internalization (0 min) in control embryos (B) and *atg-9(Q235*)* (C) mutants. Polar body contents are broken down (90 min) into a small, dim fragment in control and *atg-9(Q235*)* mutants. Scale bar is 2 μm. (D) Timing of polar body clearance in *atg-9(Q325*)* mutant embryos (n=9) compared to control embryos (n=8). Open circles denote the end of a time-lapse movie when polar body clearance did not occur. Data are presented as mean ± std dev. One-tailed t-test: n.s.= not significant p>0.1.

### Statistics

Statistical significance of polar body membrane breakdown timing was determined by performing a Student’s one-tailed t-test with unequal variance. All significant p-values were adjusted using Bonferroni corrections for multiple comparisons. Data is represented as mean ± standard deviation. Statistical significance of percent LGG-2 colocalization with the second polar body was determined by performing a one-tailed Fisher’s exact test.

### Data Exclusions

Images in which the polar body was out of view or time lapse series where embryonic development was arrested or significantly delayed were excluded. LGG-2 images that included a polar body with condensed chromosomes (before corpse membrane breakdown) were excluded from the colocalization analysis in Fig 5G. Polar bodies internalized before the 3-cell stage were excluded from the internalization analysis in Fig S3.

## Results and Discussion

To test whether ATG-9 and the ATG-16 paralogs are required for autophagosome biogenesis in early *C. elegans* embryos, we crossed a reporter for the LC3 homolog mCh::LGG-2 into *atg-9, atg-16.1,* or *atg-16.2* mutants with premature stop codons (Fig S1) (22–24). In control embryos, we observed an average of 27±15 mCh::LGG-2 puncta between the 8- and 15-cell stage (Fig 2A and F). There was a significant loss of autophagosome puncta in *atg-9(Q235*)* or *atg-16.2(W253*)* mutant embryos (Fig 2B and D), averaging 1±1 mCh::LGG-2 puncta (Fig 2F). Instead, mCh::LGG-2 weakly accumulated in nuclei and on centrosomes during early phases of mitosis in *atg-9* or *atg-16.2* mutants (Fig 2B and D), consistent with previous observations of centrosomal Atg8/LC3 proteins prior to macroautophagy induction (25, 26). These results confirm that ATG-9 is required for autophagosome formation in early embryos and that ATG-16.2 is required for LGG-2 localization to autophagosomes.

In contrast to the strong reduction in *atg-9* and *atg-16.2* mutants, we observed a milder 1.6-fold reduction in autophagosome puncta in *atg-16.1(Q356*)* mutants (Fig 2C), averaging 17±9 mCh::LGG-2 puncta per embryo (Fig 2F). However, *atg-16.2* mRNA is more highly expressed in the AB cell lineage than *atg-16.1* mRNA (27), and AB blastomeres normally engulf the second polar body (2), suggesting that ATG-16.2 is more abundant in engulfing cells and plays a larger role in LGG-2 localization to autophagosomes than ATG-16.1. Disrupting both ATG-16 homologs did not further decrease the number of autophagosome puncta, as the number of mCh::LGG-2 puncta in *atg-16.2(W253*)*; *atg-16.1(Q356*)* double mutant embryos (1±2, Fig 2E-F) did not significantly differ from *atg-16.2* single mutant embryos (1±1, Fig 2F, p>0.1). These data demonstrate that ATG-16.1 can promote LGG-2 localization to autophagosomes, consistent with results later in embryogenesis (15, 23).

As the *atg-16.2* nonsense mutants had a premature stop codon in their WD40 domain (Fig S1), we had predicted that this mutation would only disrupt non-canonical autophagy, similar to findings in mice (13, 14). However, we observed a loss of the autophagosomes that clear sperm components in the early embryo (28, 29). As premature stop codons can lead to nonsense-mediated decay (30), we asked whether *atg-16.2* mRNA levels were altered by the premature stop codon in *atg-16.2*. Semi-quantitative RT-PCR demonstrated that *atg-16.2* mRNA levels were significantly reduced in the *atg-16.2(W253*)* mutant (Fig S2), revealing that the loss of autophagosomes is likely due to reduced *atg-16.2* transcripts and consequently ATG-16.2 protein levels.

To examine whether the rare remaining autophagosome puncta in *atg-9* and *atg-16.2* mutants were capable of fusing with the polar body phagolysosome, we crossed a GFP::H2B reporter into the mutant strains so that we could observe the second polar body chromosomes after phagocytosis. In the subset of mutant embryos with one or more mCh::LGG-2 puncta, puncta were not observed in the same cell as the second polar body in 92% of *atg-9* (n=41) or 88% of *atg-16.2* (n=40) mutant embryos, which would preclude an autophagosome from fusing with the phagolysosome. In combination with the embryos that lacked any mCh::LGG-2 puncta, autophagosomes were only present in the same cell as the polar body in <8% of *atg-9* or *atg-16.2(W253*)* mutant embryos, allowing us to use *atg-9* and *atg-16.2* mutants to test the role of autophagosomes in polar body clearance.

To test whether macroautophagy contributes to phagocytic clearance of the second polar body, we first tested whether engulfment was normal in the absence of autophagosomes. A previous study had shown that LGG-1 promoted exposure of an engulfment signal on apoptotic cells, namely phosphatidylserine externalization (31), which has also been observed on dying polar bodies (2). We crossed mutants lacking autophagosomes to mCh::H2B reporters to label the polar body chromosomes and GFP::PH reporters to label the plasma membrane of the engulfing cell. In time-lapse series from control embryos, polar bodies were internalized 5±2 minutes after the 4-cell stage (Fig 1A-B, S3), consistent with previous reports (2). We did not observe a significant delay in polar body uptake in *atg-9* mutants or in *atg-16.2; atg-16.1* double mutants (Fig S3). These data show that phagocytic engulfment of polar body corpses occurs without autophagosomes.

We next tested whether autophagosomes contribute to the degradation of protein cargo within the polar body, as *atg-7* depletion to reduce Atg8/LC3 lipidation delayed cargo clearance by over an hour (2). We used the disappearance of an mCherry-tagged histone reporter in the polar body as a readout for protein cargo degradation (2, 21). In control embryos, histone reporters disappeared 86±19 minutes after internalization (Fig 3A-B and D), similar to previous results (2). When macroautophagy was disrupted in *atg-9* mutants, the disappearance timing was not significantly different from control embryos (93±12 min, p> 0.1, Fig 3C and D). These data suggest that polar body clearance occurs independent of macroautophagy, similar to previous results with *epg-8* deletion mutants (2).

We next asked what happens to the timing of polar body membrane breakdown in the absence of autophagosomes, as *atg-7* depletion to disrupt Atg8/LC3 lipidation delayed membrane breakdown after internalization almost two-fold (2). We used dispersal of the condensed histones within the phagolysosome as a proxy for membrane breakdown (Fig 1C-D, 4A), which has been shown to occur one minute after breakdown of the polar body plasma membrane (2, 21). In control embryos, histone reporters dispersed 10±3 minutes after internalization (Fig 4B-C, 4J-K). When macroautophagy was disrupted in *atg-9* or *atg-16.2* mutants, membrane breakdown timing was not significantly different from control embryos (Fig 4D-E, p>0.3). Histones dispersed 12±6 minutes after internalization in *atg-9* mutants and 10±4 min in *atg-16.2* mutants (Fig 4J). These data suggest that polar body membrane breakdown occurs independent of macroautophagy, similar to previous results with *epg-8* deletion mutants (2).

**Fig 4.**
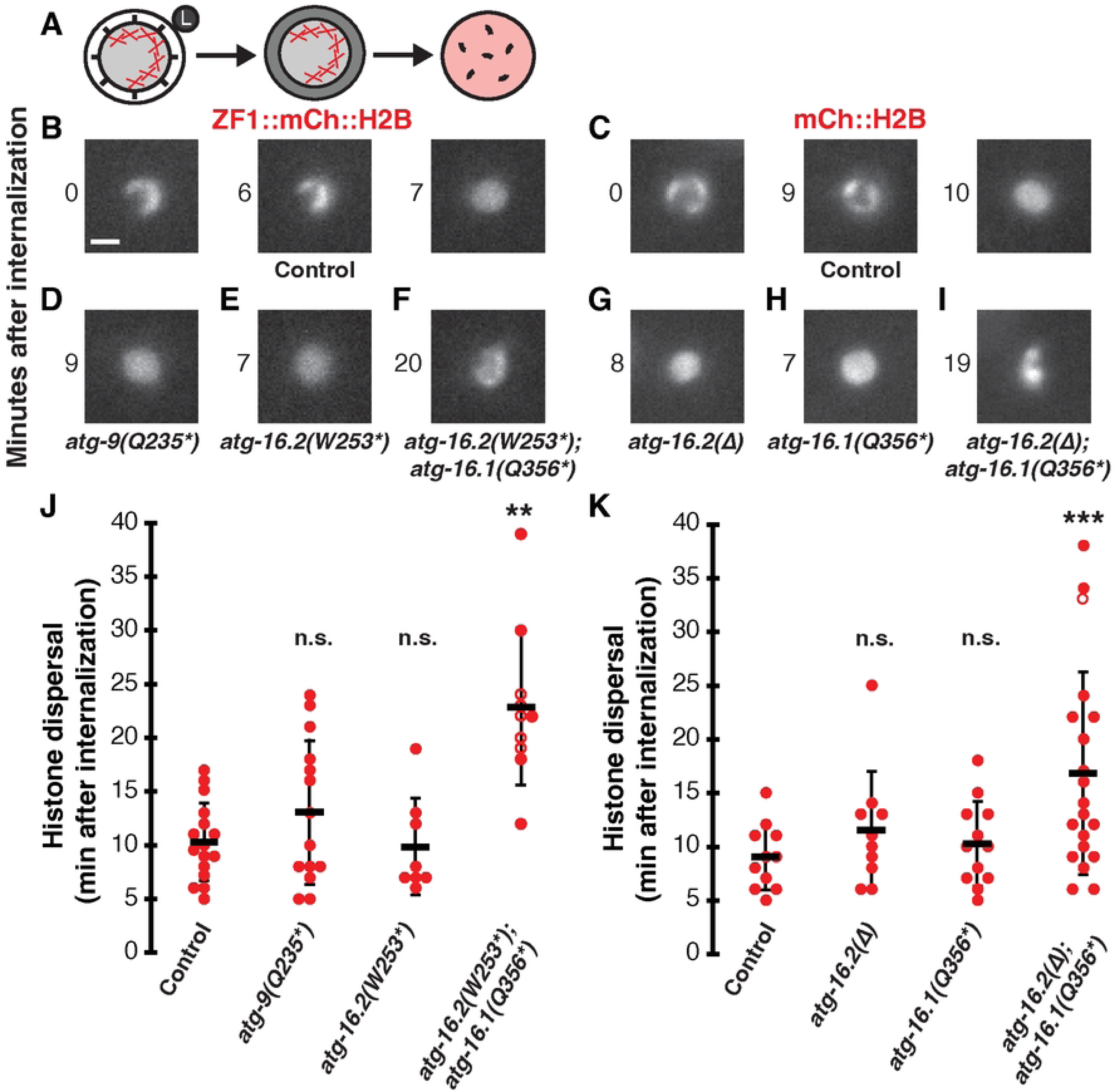
Macroautophagy does not promote breakdown of the polar body membrane, while ATG-16.1 and ATG-16.2 redundantly promote breakdown. (A) Schematic showing a polar body phagosome with condensed chromosomes (Xs) fusing with a lysosome (left), subsequent polar body membrane breakdown (middle), followed by chromosome dispersal within the phagolysosome (right). (B-C) Chromosomes in control embryos expressing ZF1::mCh::H2B (B) or mCh::H2B (C) are condensed in polar bodies at internalization (0 min) but histones disperse throughout the phagolysosome after membrane breakdown (right panel). Scale bar is 2 μm. (D-E) Histone dispersal after membrane breakdown occurs with normal timing in *atg-9(Q235*)* (D) or *atg-16.2(W253*)* (E) mutants. (F) Histone dispersal is delayed in *atg-16.2(W253*); atg-16.1(Q356*)* double mutants and chromosome condensation persists. (G-H) Histone dispersal after membrane breakdown occurs with normal timing in *atg-16.2(ok3224)* deletion mutants (G) or *atg-16.1(Q356*)* (H) mutants. (I) Histone dispersal is delayed in *atg-16.2(ok3224); atg-16.1(Q356*)* double mutants and chromosome condensation persists. (J) Timing of histone dispersal as a proxy for membrane breakdown in control ZF1::mCh::H2B embryos (n=11), *atg-9(Q235*)* (n=10), *atg-16.2(W253*)* (n=12), or *atg-16.2(W253*); atg-16.1(Q356*) (*n=20) mutants. (K) Timing of histone dispersal as a proxy for membrane breakdown in control mCh::H2B embryos (n=16), *atg-16.2(Δ)* (n=14), *atg-16.1(Q356*)* (n=8), or *atg-16.2(Δ); atg-16.1(Q356*)* (n=10) mutants. Open circles denote the end of a time-lapse movie when histone dispersal did not occur. Data are presented as mean ± std dev. Mutants were compared to controls using a one-tailed t-test with a Bonferroni correction for 3 comparisons: **p<0.01, ***p<0.001, n.s.= not significant p>0.2.

As the *atg-16.2(W253*)* allele only partially depletes *atg-16.2* mRNA levels (Fig S2), we wanted to confirm that a null allele of *atg-16.2* also had no effect on membrane breakdown. We used the *atg-16.2(ok3224)* mutant in which amino acids 125 to 299 were deleted (Δ) and the reading frame is shifted to disrupt both the CCD domain required for canonical autophagy and the WD40 domain required for non-canonical autophagy (Fig S1) (15). Deleting *atg-16.2* did not significantly delay breakdown of the polar body membrane inside the phagolysosome (p>0.2), with histone dispersal occurring 13±7 minutes after internalization (Fig 4G and K). This further confirms our findings with the nonsense allele of ATG-16.2 that polar body membrane breakdown is not promoted by macroautophagy.

We then wanted to confirm that membrane association of lipidated Atg8/LC3 promotes corpse membrane breakdown by disrupting both ATG-16.1 and ATG-16.2. In *atg-16.1 atg-16.2* double nonsense mutants, we found a significant delay in polar body membrane breakdown, with histone dispersal taking 17±9 minutes after internalization (Fig 4F and J, p<0.004). This almost two-fold delay is similar to previous results with *atg-7* knockdown (2). In double mutants using the *atg-16.2* deletion allele, we also found a significant over two-fold delay in membrane breakdown (Fig 4I and K, p<0.0005), with histone dispersal occurring 23±7 minutes after internalization. However, the balanced deletion strain rapidly went sterile and was lost, limiting our data collection. These findings confirm the role of Atg8/LC3 membrane association in promoting timely corpse membrane breakdown.

In contrast to the double mutants, we did not observe a significant delay in polar body membrane breakdown in *atg-16.1* single mutants (p>0.5), with histone dispersal occurring 10±4 minutes after internalization (Fig 4H and K), demonstrating that either ATG-16.1 or ATG-16.2 is sufficient to promote timely membrane breakdown. This suggests that ATG-16.1 and ATG-16.2 redundantly regulate Atg8/LC3 association on membranes to promote timely breakdown of the polar body membrane, while ATG-16.2 preferentially regulates Atg8/LC3 recruitment to autophagosome membranes. As autophagosomes were absent in *atg-16.2* single mutants (Fig 2D and F) and *atg-16.2* single mutants did not show a delay in membrane breakdown (Fig 4G and K), these data suggest that Atg8/LC3 lipidation on non-autophagosomal membranes promotes timely breakdown of the polar body membrane inside phagolysosomes.

To determine whether mCh::LGG-2 localized inside the polar body phagolysosome in the absence of autophagosomes, we compared the phagolysosome fluorescence to neighboring cytoplasm. As shown previously (2), mCh::LGG-2 is not visibly enriched inside polar bodies prior to membrane breakdown (Fig 5A). After membrane breakdown, mCh::LGG-2 was weakly enriched in the phagolysosome in 87% of control embryos between the 8- and 15-cell stage (Fig 5B, n=39), consistent with previous results (2). Neither *atg-9* (92%, n=36), *atg-16.1* (89%, n=35), nor *atg-16.2* (79%, n=38) mutants showed a significant difference in the percentage of embryos with polar body phagolysosomes enriched in mCh::LGG-2 between the 8- and 15-cell stage (Fig 5C-E, p>0.4). Furthermore, the fluorescence intensity of mCh::LGG-2 in the phagolysosome was not significantly different between controls and *atg-9*, *atg-16.1,* or *atg-16.2* single mutants (Fig 5G, p>0.2). Together, these data demonstrate that autophagosomes are not required for mCh::LGG-2 localization inside polar body phagolysosomes, in contrast to apoptotic phagolysosomes mid-embryogenesis (8). Therefore, it is unlikely that autophagosome fusion with phagolysosomes significantly contributes to the localization of LGG-2 in the polar body phagolysosome.

**Fig 5.**
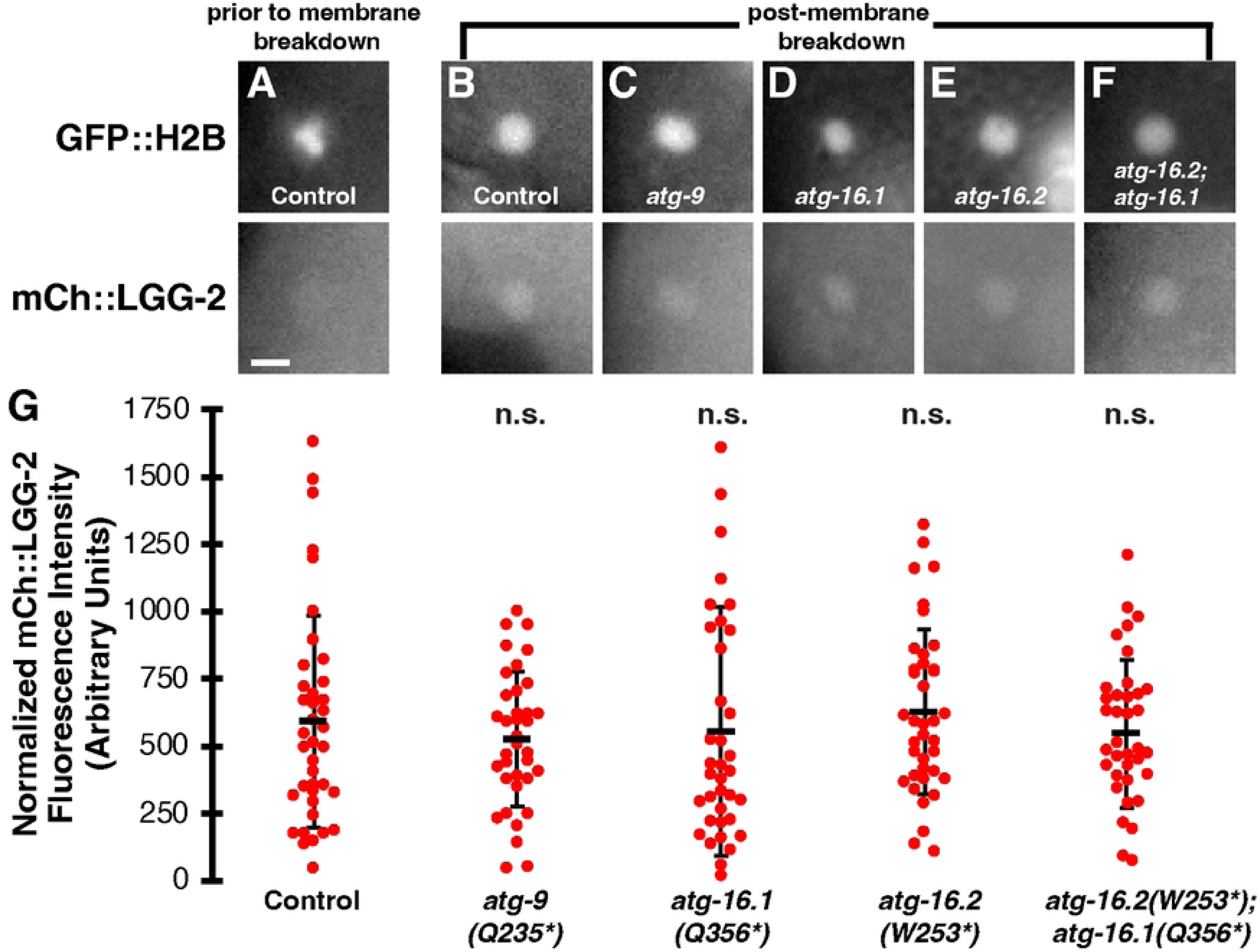
LC3 localizes in polar body phagolysosomes without autophagosomes or ATG-16 proteins. (A) Condensed polar body histones labeled with GFP::H2B are not colocalized with mCh::LGG-2 prior to corpse membrane breakdown in a control 8-cell embryo. Scalebar is 2 µm. (B-F) After polar body membrane breakdown, dispersed GFP::H2B is colocalized with mCh::LGG-2 from a control 12-cell embryo (B), an *atg-9(Q325*)* 12-cell embryo (C), an *atg-16.1(Q356*)* 8-cell embryo (D), an *atg-16.2(W253*)* 12-cell embryo (E), or an *atg-16.2(W253*); atg-16.1(Q356*)* 12-cell embryo (F). (G) There was no significant difference in the intensity of mCh::LGG-2 within the phagolysosome after polar body membrane breakdown between control and autophagosome mutant 8- to 15-cell embryos (n=39 control, 36 *atg-9*, 38 *atg-16.1*, 38 *atg-16.2,* 37 *atg-16.2; atg-16.1* embryos). Mutants were compared to controls using a one-tailed t-test: n.s.= not significant p>0.2.

We then examined whether ATG-16.1 and ATG-16.2 were redundantly required for the accumulation of LGG-2 in the polar body phagolysosome, given their redundant requirement for corpse membrane breakdown. mCh::LGG-2 was weakly enriched in the phagolysosome in 95% of *atg-16.2*; *atg-16.1* double mutant embryos between the 8- and 15-cell stage (Fig 5F, n=37), similar to control embryos (Fig 5B, p>0.2). Furthermore, the fluorescence intensity of mCh::LGG-2 in the phagolysosome was not significantly different between controls and *atg-16.2*; *atg-16.1* double mutants (Fig 5G, p>0.2). These data suggest that the mCh::LGG-2 localization observed inside polar body phagolysosomes corresponds to a non-membrane-associated pool, leaving it unclear where lipidated mCh::LGG-2 localizes to promote polar body membrane breakdown.

As LGG-2 primarily localizes in nuclei entering mitosis in autophagy and Atg8/LC3 membrane association mutants (Fig 2B-E), we examined the timing of LGG-2 accumulation in the nuclei of embryonic blastomeres. mCh::LGG-2 fluorescence increased in AB nuclei at prometaphase in control embryos (Fig 6A, Video S1), when the nuclear envelope first opens to the cytosol (32). mCh::LGG-2 fluorescence then disappeared from nuclei after anaphase (Fig 6C, Video S1), when nuclear envelope breakdown is complete. The mCh::LGG-2 fluorescence increased similarly in nuclei at prometaphase in *atg-16.2; atg-16.1* double mutant embryos (Fig 6B and D), indicating that the nuclear accumulation is independent of membrane association. These findings correlate the appearance of soluble mCh::LGG-2 with opening of both the nuclear envelope in the cytosol and the polar body plasma membrane within the phagolysosome lumen.

**Fig 6.**
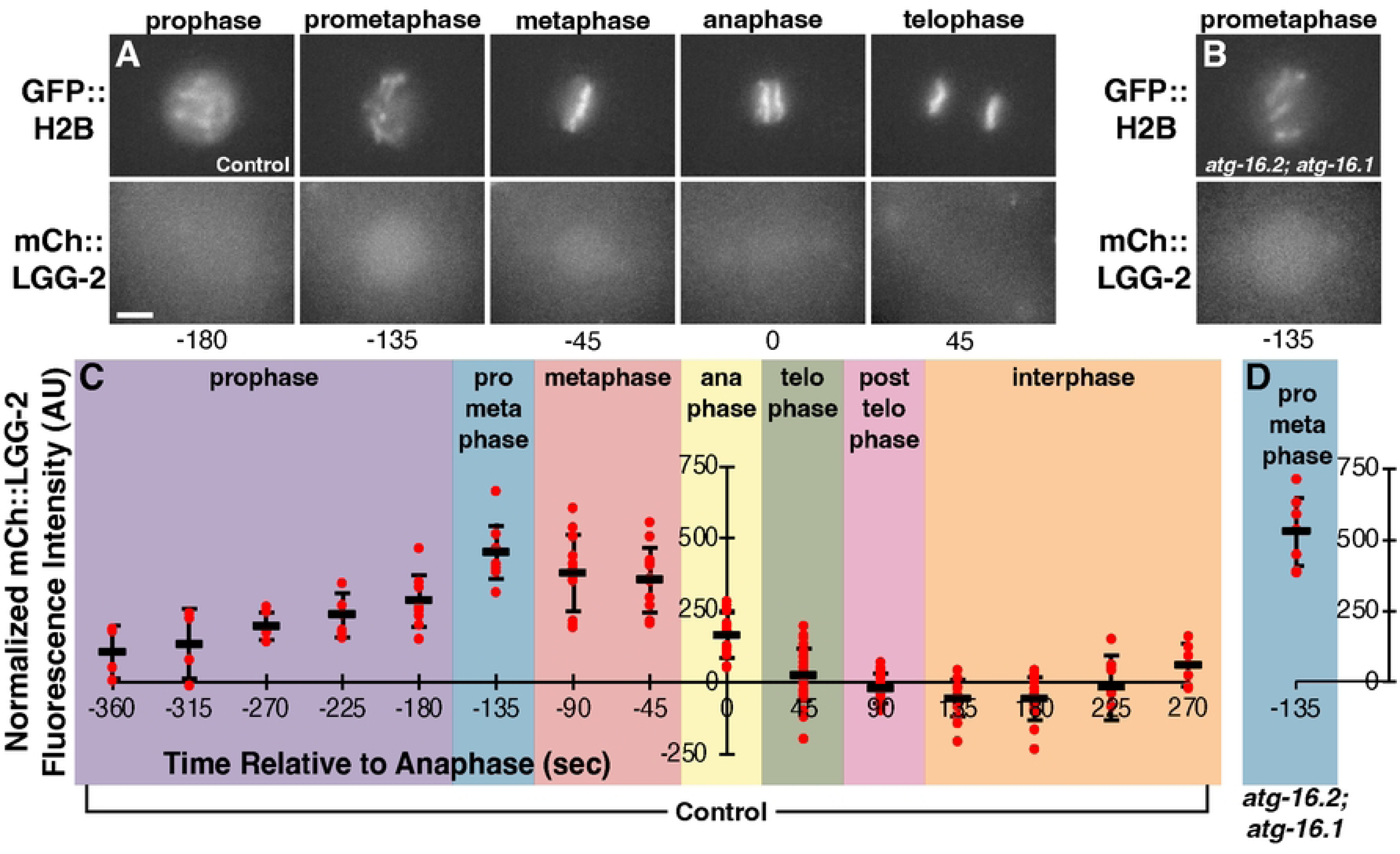
LGG-2 accumulates in mitotic nuclei as the nuclear envelope breaks down. (A-B) The mitotic nucleus of an AB cell from a control embryo (A) or *atg-16.2(W253*); atg-16.1(Q356*)* mutant embryo (B) labeled with GFP::H2B colocalizes with the LC3 reporter mCh::LGG-2 from prophase to anaphase, the semi-open phases of mitosis when there are gaps in the nuclear envelope. mCh::LGG-2 also colocalizes with centrosomes (visible in anaphase and telophase images). Scale bar is 5 µm. See also Video S1. (C) The intensity of mCh::LGG-2 on mitotic AB and P1 nuclei relative to the cytoplasm of control embryos during the transition from 2-cell to 4-cell stage measured every 45 seconds and plotted relative to anaphase timing. (D) The intensity of mCh::LGG-2 on mitotic AB and P1 nuclei relative to the cytoplasm of *atg-16.2(W253*); atg-16.1(Q356*)* mutant embryos at 135 seconds prior to anaphase is not significantly different than in control embryos (one-tailed t-test: p>0.1).

### Video S1. LGG-2 accumulates in mitotic nuclei as the nuclear envelope breaks down

A 2-cell *C. elegans* embryo developing to the 4-cell stage shows the timing of LC3 reporter mCh::LGG-2 accumulation inside the nucleus, as well as to centrosomes and spindle microtubules. Mitotic stage is visible in the merged image with histone GFP::H2B reporter. A z-series was recorded every 45 seconds, six 1.5 µm z-step images were max projected, and the projections are displayed at 5 fps using Imaris.

To determine whether the LGG-2 observed within polar body phagolysosomes was from the dying or engulfing cell, we created a degron-tagged LGG-2 reporter as a selective labeling approach. Degron tagging allows for the specific labeling of proteins protected by membranes as cytosolic degron-tagged proteins will be targeted for proteasomal degradation (18). As LGG-2 is found on both the cytosolic face of the outer autophagosome membrane and the luminal face of the inner membrane of autophagosomes, we expected a loss of LGG-2 from only the cytosolic face of autophagosomes that have sealed prior to the onset of degradation. To initiate degradation in embryonic blastomeres prior to polar body uptake at the 4-cell stage, we tagged the mCh::LGG-2 reporter with the C-terminal phosphodegrons (CTPD) from OMA-1, which are phosphorylated during the first cell division in *C. elegans* embryos. Phosphorylation leads to the degradation of CTPD-tagged proteins during the 2- and 3-cell stages (18, 33), but CTPD-mediated degradation does not occur inside polar bodies (18).

To ask whether the N-terminal CTPD tag disrupted the localization of LGG-2, we first examined CTPD::mCh::LGG-2 localization before the onset of CTPD-mediated degradation. CTPD::mCh::LGG-2 localized to discrete puncta in 1-cell embryos (Fig 7A), consistent with autophagosomal localization. To test whether the CTPD tag was able to promote degradation of autophagosomal LGG-2, we examined CTPD::mCh::LGG-2 localization over 15 minutes after CTPD-mediated degradation (Fig 7B). We found a significant reduction in the number of CTPD::mCh::LGG-2 puncta in 8- to 15-cell embryos after CTPD-mediated degradation (12±7, Fig 7C) in comparison to the mCh::LGG-2 reporter without the degron tag (27±15, Fig 2A, p<0.001) or 1-cell CTPD::mCh::LGG-2 embryos prior to CTPD-mediated degradation (19±7, Fig 7C, p<0.002). Additionally, we did not observe CTPD::mCh::LGG-2 on centrosomes and found reduced labeling of mitotic nuclei after the first cell division, suggesting degradation of microtubule-associated and nuclear LGG-2 in embryonic blastomeres. These results reveal that the degron partially reduced the labeling of autophagosomes and degraded cytosolic pools of LGG-2 in the engulfing cells.

**Fig 7.**
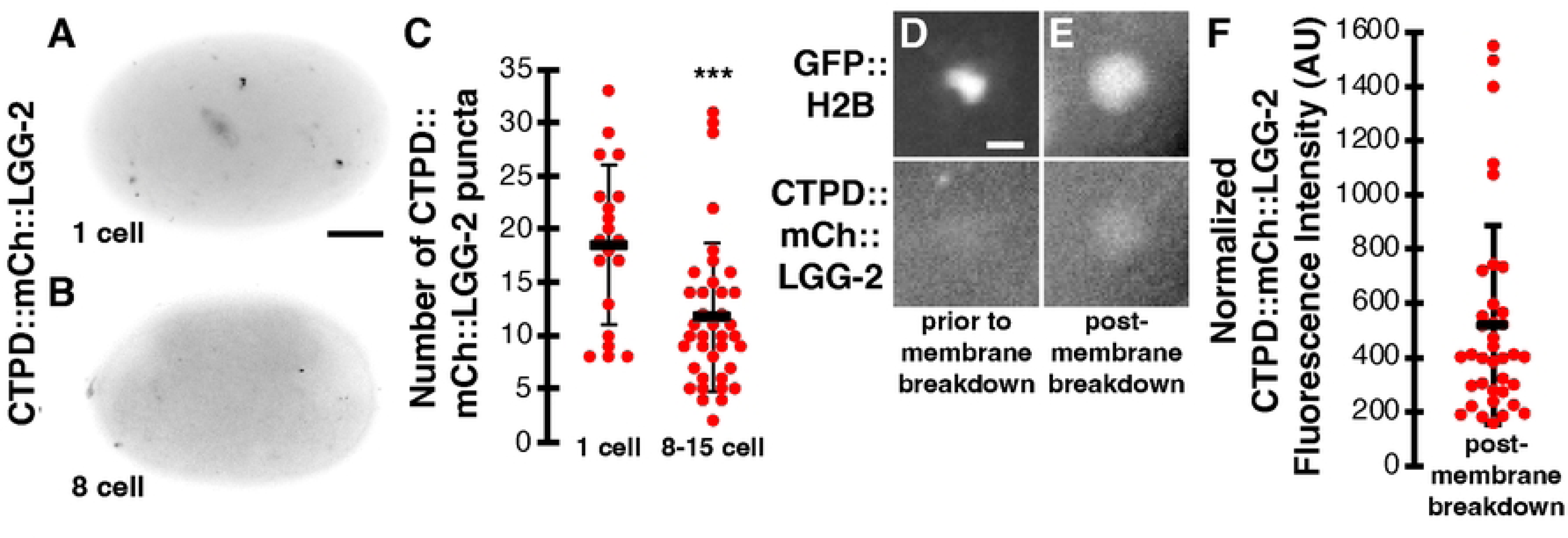
Degrading LGG-2 from engulfing cells reduces autophagosome puncta but does not disrupt LGG-2 enrichment in polar bodies. (A-B) Inverted images of control embryos expressing CTPD::mCh::LGG-2 at the indicated stage. Scalebar is 10 µm. (C) 1-cell control embryos averaged 19±7 CTPD::mCh::LGG-2 puncta per embryo (n=20), while 8- to 15-cell control embryos averaged significantly fewer: 12±7 CTPD::mCh::LGG-2 puncta (n=38). One-tailed t-test: ***p<0.001. (D) CTPD::mCh::LGG-2 is not observed within the polar body prior to corpse membrane breakdown, labeled with condensed GFP::H2B. Scalebar is 2 µm. (E) CTPD::mCh::LGG-2 is observed within the polar body post-membrane breakdown, labeled with dispersed GFP::H2B. (F) The intensity of CTPD::mCh::LGG-2 inside the phagolysosome after polar body membrane breakdown relative to the cytoplasm of 8- to 15-cell embryos (n=37). Data are presented as mean ± std dev.

Finally, we examined whether the degron-tagged mCh::LGG-2 reporter localized inside the polar body phagolysosome. We did not observe enrichment of CTPD::mCh::LGG-2 inside the phagolysosome prior to polar body membrane breakdown (Fig 7D), but CTPD::mCh::LGG-2 localized within the phagolysosome in 100% of 8- to 15-cell embryos after polar body membrane breakdown (n=37, Fig 7E), similar to mCh::LGG-2 without the degron tag. The fluorescence intensity of CTPD::mCh::LGG-2 in the polar body after membrane breakdown was brighter than the neighboring cytoplasm (Fig 7F), similar to mCh::LGG-2 without the degron tag (Fig 5G). In summary, mCh::LGG-2 enriches inside the phagolysosome after the plasma membrane of the polar body is broken down and this localization inside the phagolysosome lumen is independent of autophagosomes (Fig 2 and 5), cytosolic LGG-2 pools in the engulfing cells (Fig 7), and membrane-association of LGG-2 by ATG-16 proteins (Fig 5). Therefore, the mCh::LGG-2 fluorescence in the polar body likely comes from the dying cell after corpse membrane breakdown, although we cannot rule out the fusion of endolysosomes containing luminal LGG-2 from the engulfing cell.

## Conclusions

Together, our data suggest that phagocytic degradation of the second polar body requires the association of lipidated LC3 with membranes that are not autophagosomes. This finding is consistent with normal residual body clearance during spermatogenesis in autophagy mutant worms (6), which suggests that autophagosomes do not contribute to the phagocytic clearance of large cell fragments by gonadal sheath cells. However, our findings in early embryonic blastomeres contrast with the role of autophagosome fusion in the degradation of apoptotic corpses mid-embryogenesis (8). One possibility is that apoptotic and non-apoptotic corpses induce different clearance mechanisms. However, polar bodies and residual bodies expose phosphatidylserine and depend on the same signaling pathways as apoptotic cells for their uptake (2, 6). In addition, polar body and residual body phagosomes mature using similar Rab GTPase pathways as apoptotic phagosomes, leaving it unclear whether different phagocytic cargos have different needs for autophagosomes.

Alternatively, the difference in the role of autophagy between polar body and apoptotic corpse clearance may lie in the engulfing cells. Early embryonic blastomeres and mid-embryonic differentiated cells may have different metabolic states or types of autophagosomes and autolysosomes. The early embryonic cells that engulf the polar body have been isolated in an eggshell for <2 hours, while the apoptotic corpse phagosomes fused with autophagosomes after >5 hours of isolation inside the eggshell (8). The additional time without external nutrients could lead to starvation-induced autophagy, consistent with the abundance of autophagosomes and autolysosomes at mid-embryonic stages. Indeed, ∼40% of autophagosomes fusing with apoptotic phagolysosomes contained a lysosomal nuclease, indicating that autolysosomes also fuse with phagolysosomes (8). In contrast, early embryonic cells contain several cargos that are specifically targeted for autophagic degradation, including sperm components degraded by allophagy and RNA granules degraded by aggrephagy (28, 29, 34). Therefore, what distinguishes the autophagosome dependence of phagocytic clearance may be the metabolic state of the cell or the prevalence of lysosomes containing autophagic cargo.

Further research is needed to determine how lipidated Atg8/LC3 family proteins promote polar body clearance, including which membrane ATG-16 proteins localize lipidated Atg8/LC3 to for corpse membrane breakdown. Although Atg8/LC3 lipidation factors promote breakdown of the polar body membrane, we rarely observed LGG-1 or LGG-2 localization to the phagosome prior to membrane breakdown with fluorescent protein-tagged reporters (2). Furthermore, all LGG-1 and LGG-2 reporters examined showed colocalization of Atg8/LC3 proteins with histone markers after corpse membrane breakdown rather than the surface labeling of phagosomes predicted for LC3-associated phagocytosis (This study and (2). We also observed dynamics for mCh::LGG-2 independent of membrane association by ATG-16 proteins (Fig 5-6), but ATG-16 proteins promote timely corpse membrane breakdown (Fig 4). One possibility is that the N-terminal fluorescent protein tags on the Atg8/LC3 reporters disrupt Atg8/LC3 association to phagosome membranes but not autophagosome membranes, consistent with ATG16L1 interacting with different proteins while localizing Atg8/LC3 proteins during autophagy or LAP/CASM (13, 14). Alternatively, as Atg8/LC3 reporters increase their fluorescence in response to nuclear membrane breakdown during mitosis as well as corpse membrane breakdown inside phagolysosomes, the observed changes in fluorescence of mCh::LGG reporters could be due to interactions of soluble Atg8/LC3 proteins between mixing organelle compartments, obscuring the membrane-associated Atg8/LC3 proteins that function in corpse clearance. Thus, it appears that a low level of membrane-associated Atg8/LC3 is sufficient to promote corpse membrane breakdown inside phagolysosomes.

## Acknowledgements

The authors would like to thank Gabriela Paredes, Riley Harrison, and Brianna Brost for technical assistance and Ahmad Fazeli and Chase White for suggestions on the manuscript. Strains were provided by Jeremy Nance, Vincent Galy, Renaud Legouis, and the CGC, which is funded by NIH Office of Research Infrastructure Programs (P40 OD010440). This work was funded by NIH grant R15 GM143727-01 (AMW).

---

**Fig S1. ATG-9 and ATG-16 alleles used in this study.**

ATG-9 has four transmembrane domains (TM) and two alpha-helices predicted to be partially embedded in the membrane (M), based on Guardia, Tan (35). Most of the protein is cytosolic, except for two small luminal domains (L). ATG-16.1 and ATG-16.2 have an N-terminal domain predicted to bind ATG-5 and ATG-12, a central coiled-coil domain (CCD) important for macroautophagy, and a C-terminal WD40 domain important for non-canonical autophagy (LAP/CASM). Positions of point mutations and the *ok3224* deletion are indicated.

**Fig S2. *atg-16.2* mRNA levels are reduced by a premature stop codon in the WD40 domain.** (A) Three biological replicates of wild-type N2 and *atg-16.2(gk145022[W253*])* cDNA were co-amplified for *atg-16.2* and *tat-5* as a loading control. (B) Graph of normalized subtracted fluorescence intensity ratios. The *atg-16.2(gk145022[W253*])* mutant band is significantly reduced compared to wild-type using a one-tailed t-test. ***p<0.001.

**Fig S3. Macroautophagy is not required for polar body internalization.**

Timing of polar body internalization after the 4-cell stage. Control embryos averaged 5±2 minutes after the 4-cell stage (n=11). There was no significant delay in internalization in *atg-9* single mutants (5±2, n=10) or *atg-16.2(W253*); atg-16.1(Q356*)* double mutants (6±3, n=10). Data are presented as mean ± std dev. One-tailed t-test, p>0.3.

**Table S1.**
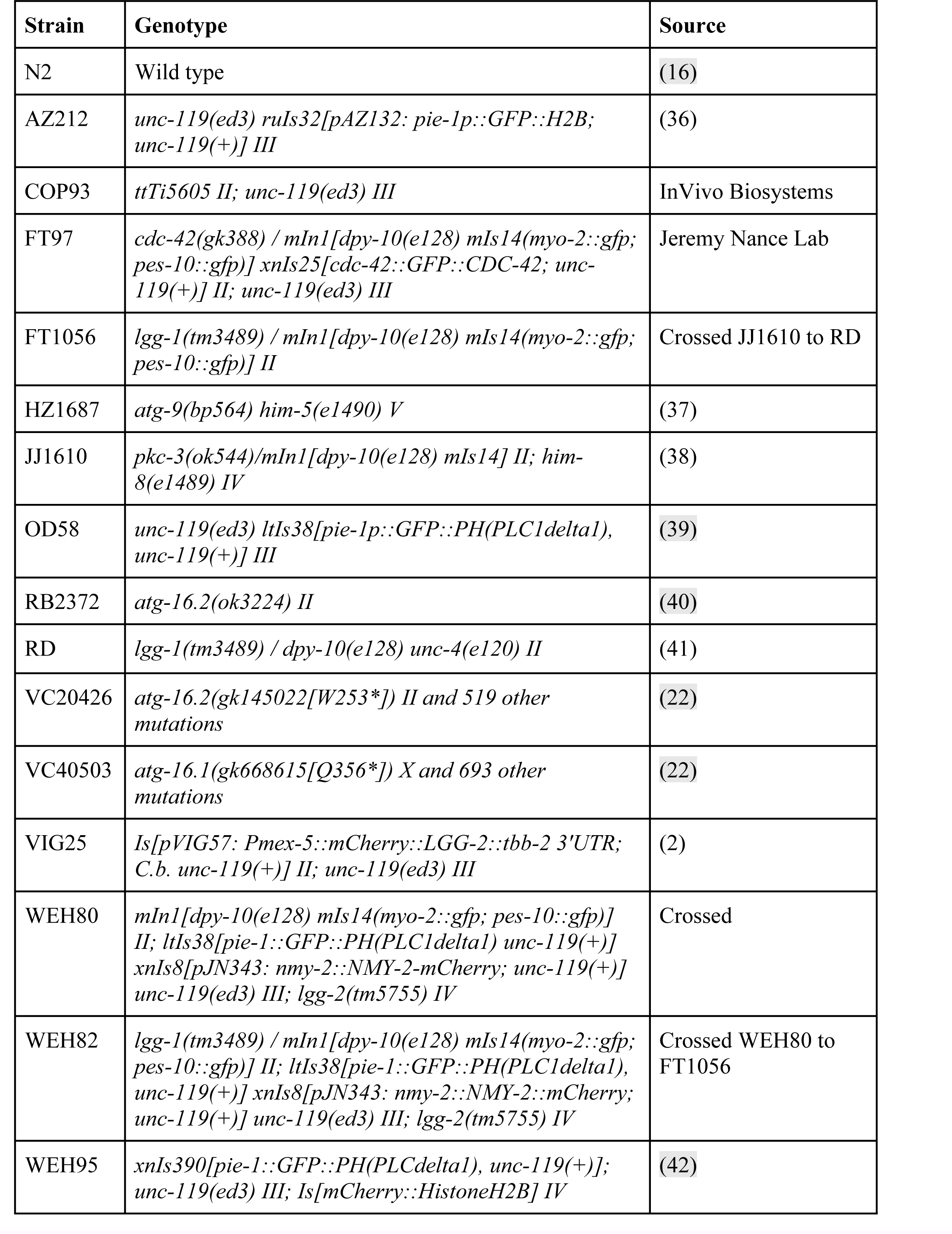
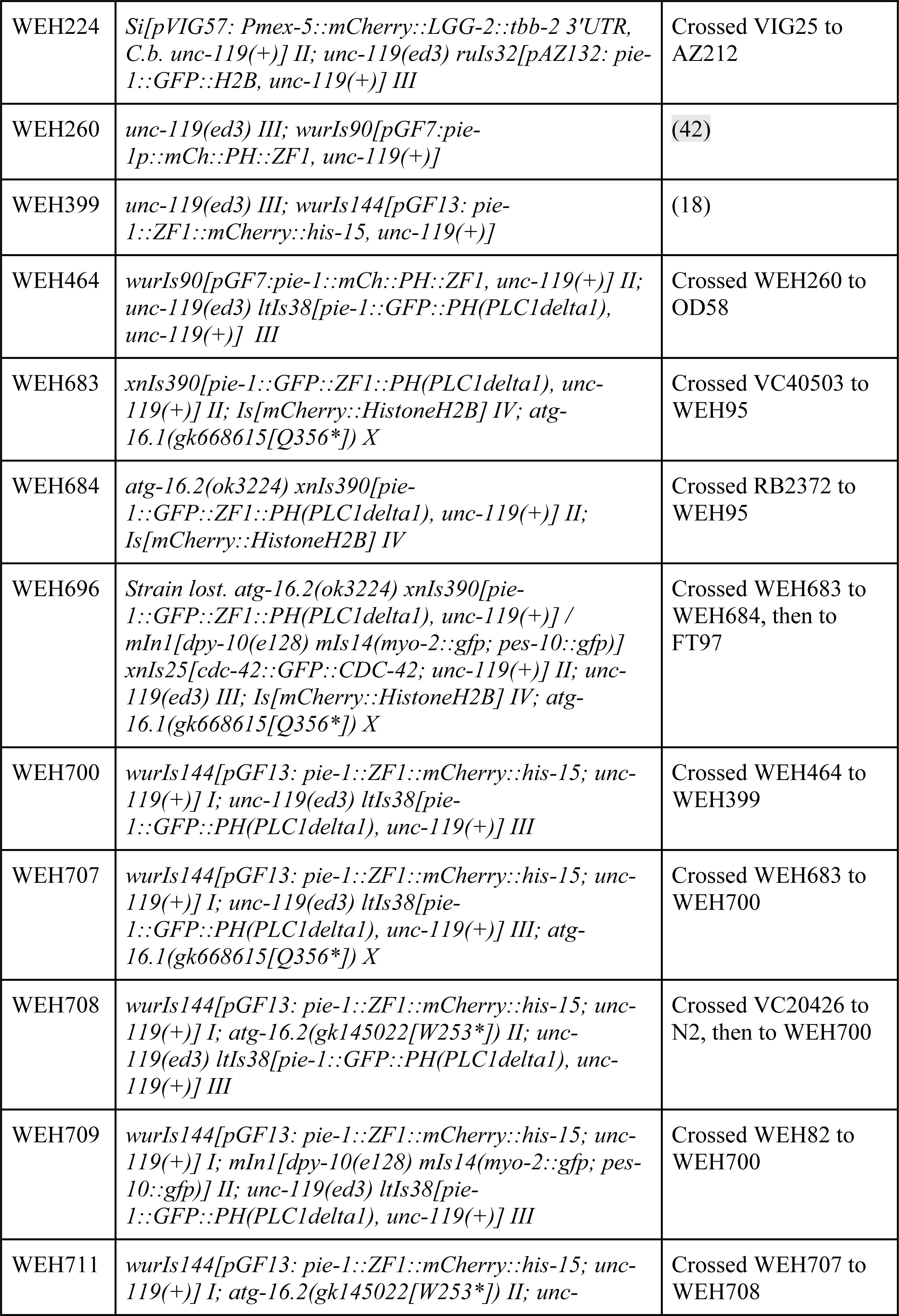
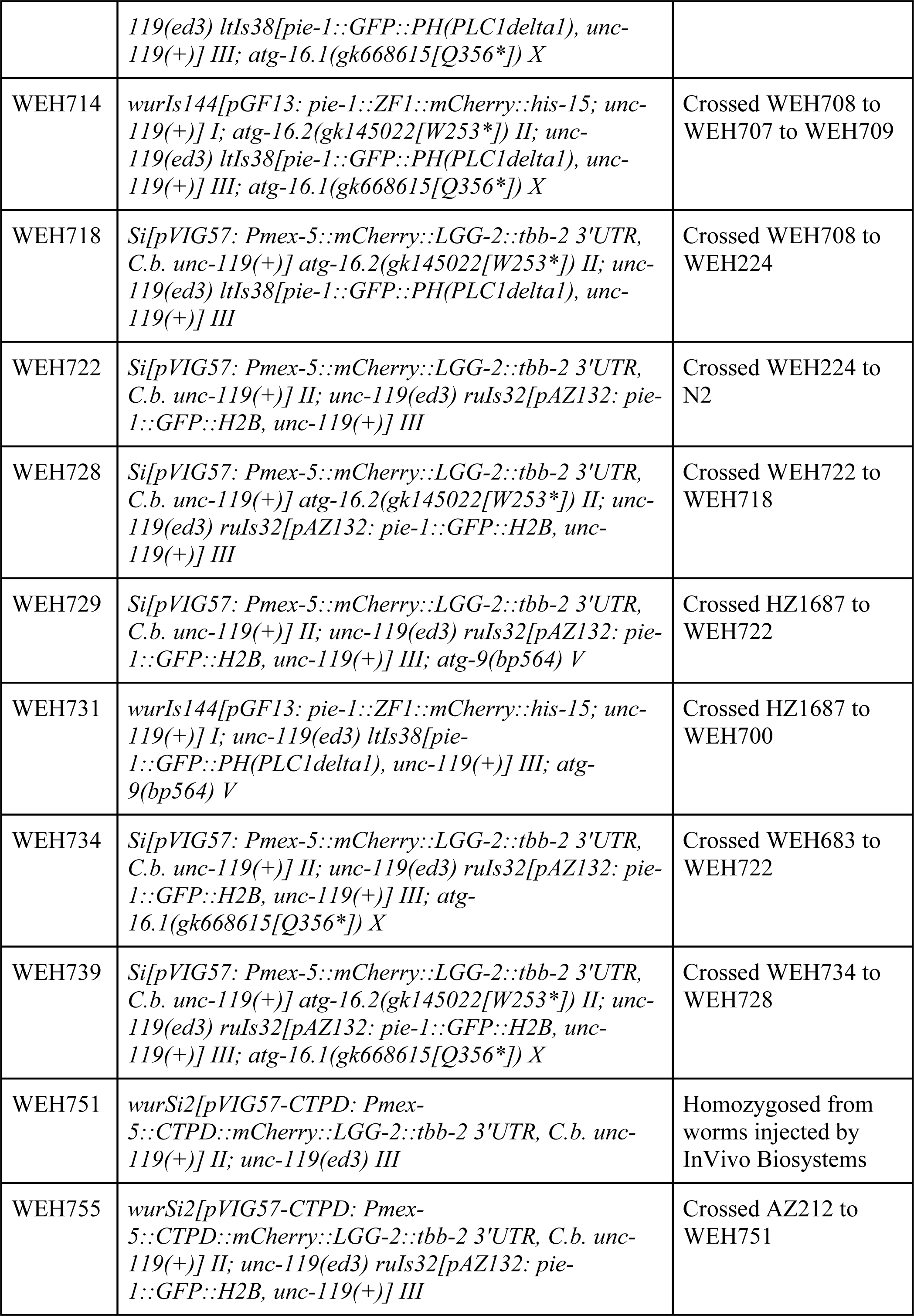
Worm strains.

**Table S2.**
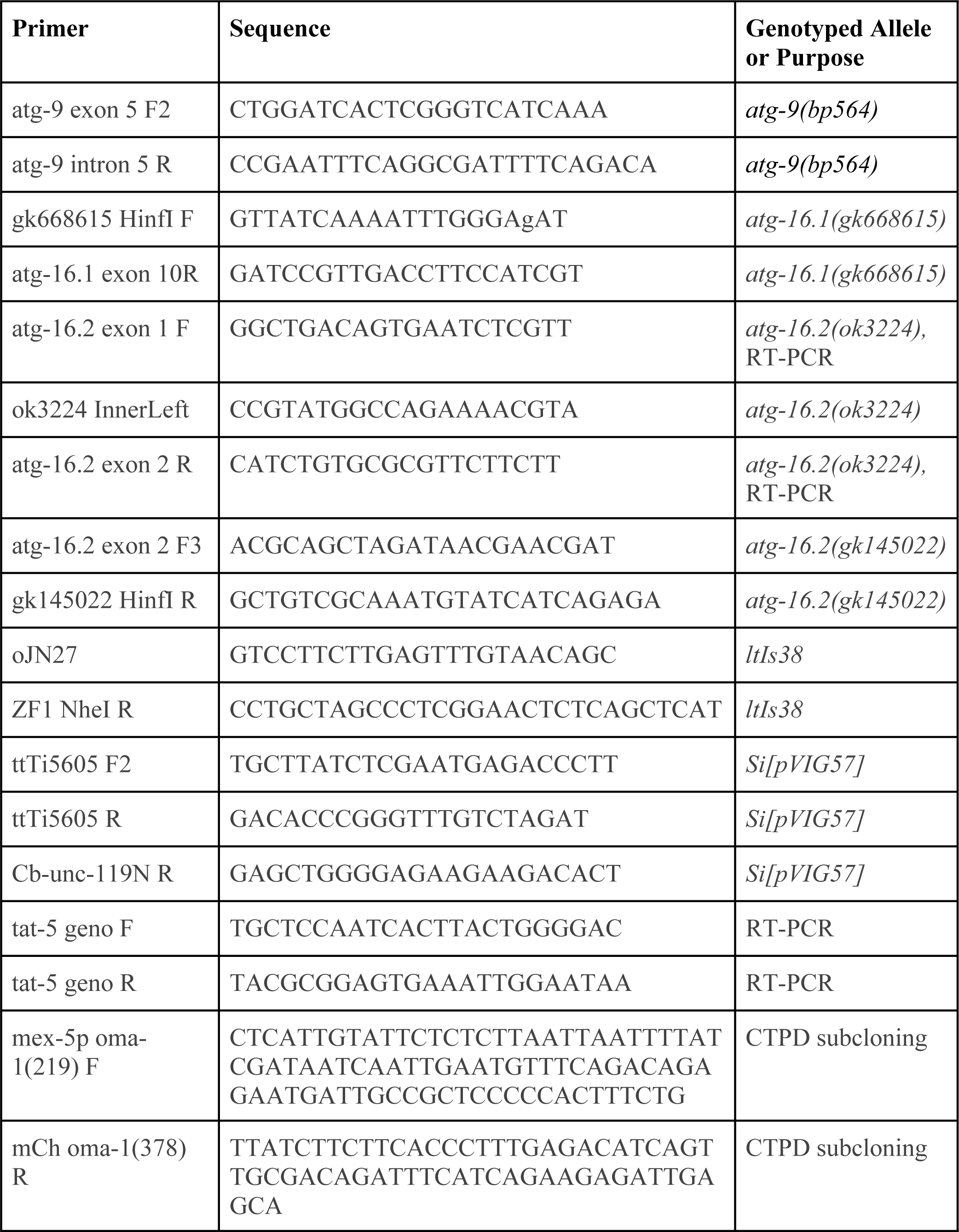
Oligonucleotide primers.

